# Leveraging immonium ions for identifying and targeting acyl-lysine modifications in proteomic datasets

**DOI:** 10.1101/2020.04.20.052191

**Authors:** John M. Muroski, Janine Y. Fu, Hong Hanh Nyugen, Rachel R. Ogorzalek Loo, Joseph A. Loo

**Author notes:** Equal contribution (John M. Muroski, Janine Y. Fu).

## Abstract

Acyl modifications vary greatly in terms of elemental composition and site of protein modification. Developing methods to identify these modifications more confidently can help assess the scope of these modifications in large proteomic datasets. Herein we analyze the utility of acyl-lysine immonium ions for identifying the modifications in proteomic datasets. We demonstrate that the cyclized immonium ion is a strong indicator of acyl-lysine presence when its rank or relative abundance compared to other ions within a spectrum is considered. Utilizing a stepped collision energy method in a shotgun experiment highlights the immonium ion strongly. Implementing an analysis that accounted for features within each MS^2^ spectra, this method allows peptides with short chain acyl-lysine modifications to be clearly identified in complex lysates. Immonium ions can also be used to validate novel acyl-modifications; in this study we report the first examples of *3-*hydroxylpimelyl-lysine modification and validate them using immonium ions. Overall these results solidify the use of the immonium ion as a marker for acyl-lysine modifications in complex proteomic datasets.

**Statement of Significance:** Acyl-lysine modifications come in a variety of elemental compositions. There is increasing evidence that these modifications can have a functional effect on protein and are present in proteomes across all domains of life. Here we describe a new method that can allow for more confident identification of acyl modifications in proteomes by utilizing the immonium ion of these modifications. Our utilization of these ions allows for more comprehensive insight into the role of acyl modifications at the systems level.

## 1. Introduction

As proteomic technology advances, the depth and breadth of proteomic data that can be analyzed has increased. This increased scope of proteomic information has resulted in a plethora of information on post-translational modifications (PTMs), which can affect protein function and modulate protein activity in a cell without the energetic burden of new protein expression.^[1–5]^ These modifications fall into different classes based on their physicochemical properties, such as phosphorylation, glycosylation, oxidation and many more. One class of modification that has been observed across biological systems is lysine acylation.^[6–8]^ Acetylation has been long known as an epigenetic regulator, modulating protein expression by modifying lysine side chains on histone tails.^[9–11]^ Later, it was discovered that acetyl and other acyl modifications not only impact histone function, but are also ubiquitous in mammalian metabolic pathways as well as in other eukaryotic and prokaryotic systems.^[6,9]^ In eukaryotic organisms, these modifications have been shown to correlate with aging as well as an organism’s metabolic state.^[7,8]^ In prokaryotes, these modifications have been shown to impact the activity of enzymes in metabolic pathways.^[12,13]^ Acylation even plays a role in viruses, as viral proteins have been shown to require acetylation to function.^[14]^

Acyl-modifications have proven to be challenging to identify on a proteome-wide scale for a number of reasons. These modifications tend to have low stoichiometries relative to unmodified proteoforms; *e.g.*, mitochondrial acylation stoichiometries are around 0.02%.^[15]^ This low abundance makes it difficult to consistently identify modified peptides in complex samples using untargeted proteomic methods, and enrichment strategies during sample preparation are often necessary.^[16]^ A second challenge is that acyl modifications come in many different varieties. Chain length, elemental composition, and degree of unsaturation differ between different types of acylations.^[4]^ For example, an acetyl group and a propionyl group differ by one carbon. Although it is often unclear whether these modifications assume different functions, identifying them requires a priori knowledge of which acyl compositions might be present. Acylation can occur non-spontaneously; that is, reactive intermediates in certain metabolic pathways may modify certain primary amines even without enzymatic catalysis.^[17]^ Such processes suggest that a given residue could be tagged by many acyl groups, even on a single peptide. Not including all possible acyl modifications during the sequence database search step in a proteomic workflow may miss critical information about a given protein, particularly when quantification is involved. Broadening the number of possible modifications considered, however, can increase the false discovery rate (FDR) if too many PTMs are considered or, if FDR is treated properly, will reduce the sensitivity for peptide identification. All of these challenges add to the difficulty of identifying the full range of possible acyl modifications.

Physicochemical differences in PTMs have meant that different experimental strategies may need to be considered to detect and localize different modifications optimally. For example, the phosphate group of phosphopeptides is often labile to collision induced dissociation (CID) and its associated mass shift may not be apparent in the MS^2^ spectra or it may migrate to another residue.^[18]^ To overcome this limitation, phosphoproteomic studies have adopted strategies that utilize electron transfer dissociation (ETD) as an alternative MS^2^ dissociation method.^[18,19]^ Glycosylated peptides also present challenges with MS fragmentation given the fragility and complexity of their structures. One strategy for identifying these modifications involves the use of collisional energy stepping in conjunction with the detection of characteristic oxonium ions associated with specific glycans.^[20]^ Oxonium ions are low mass ions resulting from fragmentation of oligosaccharides and glycopeptides that are used to identify glycan species in complex samples.^[21–23]^

Similarly, immonium ions can serve as a diagnostic marker for a specific acyl-lysine PTM.^[24,25]^ Immonium ions are internal product ions resulting from two-bond cleavages that retain a single amino acid side-chain. These ions are common in peptide tandem mass spectra and can often verify the presence of certain amino acids in a peptide.^[26,27]^ It has been shown that in addition to the canonical amino acids, immonium ions can be generated for acyl-lysine residues. The immonium ion for acetyl-lysine is observed at *m/z* 143.1179; however, a related ion is often observed at *m/z* 126.0913 that originates from cyclization.^[24,28]^ Other acyl-lysines present unique immonium ions.^[25]^ These diagnostic indicators are typically used for post-identification validation, because their presence and intensity depend heavily on sequence context and instrument parameters. Here, we posit a means to overcome limitations in the use of immonium ions by optimizing collision energies and by defining comparative criteria. This strategy can be extended to identifying novel acyl PTMs in large proteomic data sets.

## 2. Experimental Section

### 2.1 Reagents and Materials

Synthetic peptides (lyophilized, >95% purity) were obtained from Genscript, Inc. and reconstituted in water. The sequences, which originate from *Syntrophus aciditrophicus*, are: **K**STPEAMAK, F**K**DEIPVVIK, STDP**K**GPSVR, with the lysine residues indicated in bold containing an acetyl-, butyryl-, or crotonyl-modification on the lysine ε-amine.

### 2.2 Preparation and Digestion of Acyl-Bovine Serum Albumin

Acetylated-bovine serum albumin (BSA) was obtained from Promega (Product #R3961) and diluted in 100 mM ammonium bicarbonate. Butyrylated-BSA (was prepared in a process adapted from Baez, *et al.*^[29]^ Butyric anhydride (∼25 μmol) was added to 100 μL of a 1 mg/ml solution of BSA dissolved in 100 mM ammonium bicarbonate (Sigma Aldrich, Product #A8022). The solution was incubated for 20 minutes at 4°C. The pH of the solution was then adjusted to pH ∼8 using ammonium hydroxide. The process was then repeated two more times. Subsequently, hydroxylamine hydrochloride was added at 50% (*w/v*) of the final concentration and the pH was readjusted with NH_4_OH to ∼8 to reverse adventitious *O-*acylation. The solution was incubated at room temperature overnight. Butyrylated-BSA was then buffer exchanged into 100 mM ammonium bicarbonate using 10kD MWCO Amicon spin filters (Millipore).

Acylated- and non-acylated-BSA were heated to 95°C for 10 minutes, disulfide-reduced with 20 mM dithiothreitol (DTT) for 1 hour at 60°C, and alkylated with 50 mM iodoacetamide for 45 minutes at room temperature in the dark. Excess iodoacetamide was quenched with DTT and the samples were digested overnight with endoproteinase GluC (1:100) at room temperature (New England Biolabs, Product #P8100S). Digested peptides were dried in a vacuum concentrator, acidified with 0.1% acetic acid, and desalted with STAGE tips assembled from 3M Empore C18 Solid Phase Extraction Disks^[30]^ and dried again. Peptides were reconstituted in LC-MS injection buffer (3% acetonitrile, 0.1% formic acid) and quantified by Pierce Quantitative Fluorometric Peptide Assay (Thermo Scientific, Product #23290).

### 2.3 Peptide Preparation from *S. aciditrophicus* Cells

Cells were harvested from tricultures of *Syntrophus aciditrophicus, Methanosaeta concilli*, and *Methanospirillum hungatei* grown with benzoate as the carbon source. Peptides were prepared from cell pellets using enhanced filter-aided sample preparation (eFASP) as described by Erde, *et al*.^[31,32]^ Briefly, cells were lysed in 4.0% (*v/v*) ammonium lauryl sulfate, 0.1% (*w/v*) sodium deoxycholate, and 5 mM tris(2-carboxyethyl)phosphine in 100 mM ammonium bicarbonate. The lysate was exchanged into a buffer containing 8 M urea, 0.1% (*w/v*) sodium deoxycholate and 0.1% (*w/v*) *n-*octyl glucoside. Proteins were alkylated in 17 mM iodoacetamide and digested with trypsin in a buffer containing 0.1% (*w/v*) sodium deoxycholate, 0.1% (*w/v*) *n-*octyl glucoside and 100 mM ammonium bicarbonate. Peptides were digested overnight at 37°C. Tryptic peptides were desalted with STAGE tips as described earlier.

### 2.4 Mass Spectrometry (LC-MS/MS) Analysis

Processed peptides were measured by reversed phase liquid chromatography-tandem mass spectrometry (LC-MS/MS) on an EASY nLC1000 (Thermo Scientific) coupled to a quadrupole orbitrap mass spectrometer (Q-Exactive, Thermo Scientific). Peptides (100 ng of acyl-BSA) were loaded onto an Acclaim PepMap100 C18 trap column (Thermo Scientific, Product #16-494-6, 75μm x 2cm, 100 Å) and separated on an Acclaim PepMap RSLC C18 analytical column (Thermo Scientific, Product #03-251-873, 75μm x 25μm, 100Å). The LC buffers were buffer A (0.1% formic acid) and buffer B (0.1% formic acid in 100% acetonitrile), and peptides were eluted at 300nL/min. For acyl-BSA experiments, the LC gradient used was 3-35% B in 30 min, 35-50% B in 5 min, and 50-80% B in 2 min. For synthetic acylated peptides spiked into lysates, 1 nM of each acyl-peptide was added to 100 ng of a HeLa tryptic digest standard (Thermo Scientific, #PI88329) for a final ratio of 1 fmol acyl-peptide/100 ng of HeLa digest. For the triculture digests containing *S. aciditrophicus*, 200 ng were loaded. HeLa and triculture analyses used the gradient 3-20% B in 62 min, 20-30% B in 31 min, 30-50% in 5 min, and 50-80% in 2 min.

The mass spectrometer was operated in a data-dependent acquisition mode with an *m/z* 300-1800 MS scan acquired at 70,000 resolution using an automatic gain control (AGC) target of 1E6 (maximum fill: 100 ms). Collision induced dissociation MS/MS spectra were acquired at 17,500 resolution, AGC (maximum fill: 80 ms) of 1E5, and a normalized collision energy of 27 (unless otherwise indicated) on the top 10 most abundant precursor ions.

### 2.5 Proteomic Data Analysis

#### 2.5.1 Acyl-BSA Data

RAW files were converted into MGF format and peak lists were submitted to Mascot (version 2.5; Matrix Science) and searched against the BSA sequence supplemented with protein sequences of common contaminants. GluC was specified as the cleavage enzyme with up to 6 missed cleavages considered, and a precursor mass tolerance of 10 ppm and product mass error of 0.02 Da. Cysteine carbamidomethylation (+57.021464), methionine oxidation (+15.994915), and the respective acyl-lysine modification, acetyl (+42.010565) or butyryl (+70.041865) were set as variable modifications. Peptide spectral matches (PSMs) were filtered to 1% false discovery rate using the target-decoy strategy.

#### 2.5.2 Acyl-modifications in HeLa and S. aciditrophicus

All HeLa cell data was analyzed using Mascot (version 2.5). Files were searched against the UniProt Human database (obtained as of January 23, 2019) supplemented with common laboratory contaminants and the three spiked synthetic peptide sequences. The *S. aciditrophicus* triculture RAW files were processed through ProteomeDiscoverer (version 1.4), using Mascot for the database search. Files were searched against UniProt *S. aciditrophicus, Methanosaeta concilli, and Methanospirillum hungatei* sequence databases that were concatenated and supplemented with contaminant sequences (obtained as of July 8, 2019). The search parameters for both datasets were: enzyme specificity, trypsin; maximum number of missed cleavages, 2; precursor mass tolerance, 10 ppm; product mass error allowed, 0.02 Da; variable modifications included cysteine carbamidomethylation, methionine oxidation, lysine acetylation, lysine butyrylation, and lysine crotonylation (+68.026215). PSMs were filtered to 1% false discovery rate using the target-decoy strategy.

#### 2.5.3 Immonium Ion Analysis for Acylated BSA

From the Mascot search results (DAT files), PSMs with an ion score >25 were considered for further immonium ion analysis. An in-house Python script was utilized to extract MS/MS spectra corresponding to the PSMs containing the immonium ion of interest for further characterization. For all datasets, the mass tolerance was set to 10 ppm for the immonium ion of interest. Similar analysis was performed on the Svinkina *et al*.^[33]^ data set (MassIVE MSV000079068).

## 3. Results

### 3.1 The *m/z* 126 Immonium Ion is Present in Great Abundance in an Acetyl-Lysine Dataset

Previous reports have indicated that the *m/z* 126 ion (126.0913) (hereafter called the “126 ion”) is a diagnostic marker ion for acetyl-lysine.^[24]^ To determine the prevalence of the 126 ion in peptide spectral matches (PSMs), we investigated peptides from GluC-digested acetylated BSA and focused on comparing PSMs with and without acetyl-lysine, as indicated by the MASCOT search algorithm. Of those identified as acetylated 97.3%, displayed the 126 ion in MS/MS spectra (**Table 1**). However, the 126 ion was also present in 65.7% of the non-acetylated spectra. There are several possible ways that non-acetyllysine containing peptides can yield an ion with the exact mass for the 126 ion, including *“a*-ion” type products that subsequently lose NH_3_ from sequences containing Gly-Ile, Gly-Leu, or Ala-Val and the reversed sequences. To verify that this trend of observing 126 ions from non-acetylated peptides appears in more complex lysates, we performed the same analyses on a dataset from a previous report that utilized immunoprecipitation to enrich acetylated peptides in Jurkat E6-1 cells.^[33]^ This dataset showed that while 96% of acetylated PSMs had a 126 ion, 73.9% of the non-acetylated PSMs also had a 126 product ion present (**Table 1**). That the 126 ion appears in both acetylated and non-acetylated PSMs agrees with previous work in terms of the 126 ion’s sensitivity as an acetyl-lysine marker, but not in terms of its specificity.^[24]^ This may reflect the higher sensitivity of current mass spectrometers over those used in previous studies.^[24]^ It may also be a result of the previous studies’ metrics for determining false positives which relied on the presence of specific dipeptide cleavages that may not have been present in spectra with limited sequence information.^[24]^

**Table 1.**
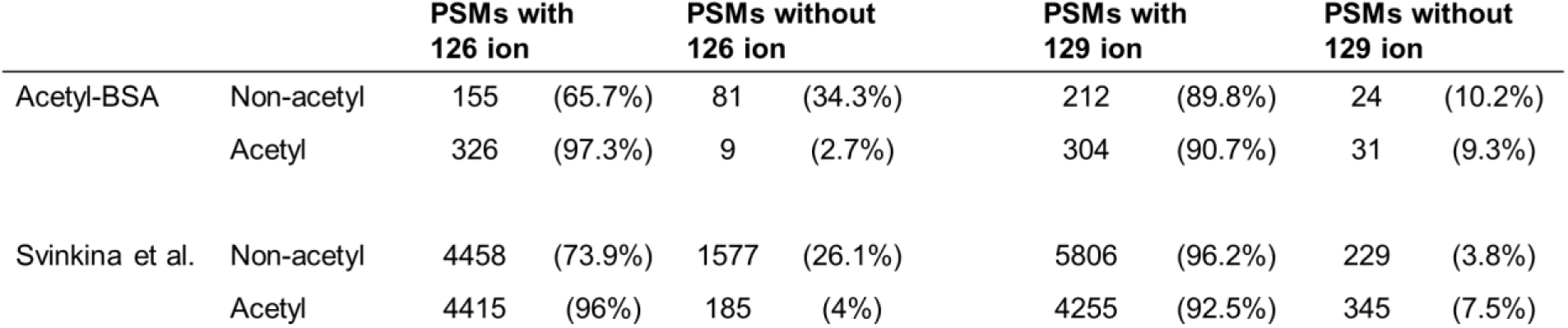
Distribution of PSMs containing 126 and 129 immonium ions from acetylated BSA (27 NCE) and from Jurkat E6-1 cells (25 NCE) (Svinkina *et al*.)^[33]^

|  |  | PSMs with<br>126 ion |  | PSMs without<br>126 ion |  | PSMs with<br>129 ion |  | PSMs without<br>129 ion |  |
| --- | --- | --- | --- | --- | --- | --- | --- | --- | --- |
| Acetyl-BSA | Non-acetyl | 155 | (65.7%) | 81 | (34.3%) | 212 | (89.8%) | 24 | (10.2%) |
|  | Acetyl | 326 | (97.3%) | 9 | (2.7%) | 304 | (90.7%) | 31 | (9.3%) |
| Svinkina et al. | Non-acetyl | 4458 | (73.9%) | 1577 | (26.1%) | 5806 | (96.2%) | 229 | (3.8%) |
|  | Acetyl | 4415 | (96%) | 185 | (4%) | 4255 | (92.5%) | 345 | (7.5%) |

### 3.2 Using the 126:129 Ion Abundance Ratio as an Acetyl-Lysine Indicator

Given the ubiquity of the 126 ion in MS^2^ spectra, a more specific diagnostic ion metric is needed to distinguish between acetylated and non-acetylated PSMs. Other low mass ions that may indicate the presence of an unmodified lysine include the *m/z* 101 (101.1079) immonium ion and a diagnostic ion at *m/z* 129 (129.1023).^[26]^ The *m/z* 101 ion was rarely observed within lysine-containing PSMs and, when present, did not clearly differentiate acetylated from non-acetylated peptides (**Supplemental Table S1**). The *m/z* 129 diagnostic ion (hereafter called the “129 ion”), however, was present more often and proved to better indicate the presence of unmodified lysine (**Figure 1**). Similar to the low specificity of the 126 ion for acetyl-lysine containing peptides, the 129 ion was not very specific for unmodified lysine-containing peptides; 89.8% and 90.7% of non-acetyl and acetyl PSMs, respectively, contained the 129 ion (**Table 1**). Similarly, the prevalence of the 129 ion in acetyl and non-acetyl PSMs was also noted by Svinkina, *et al.*^[33]^

**Figure 1.**
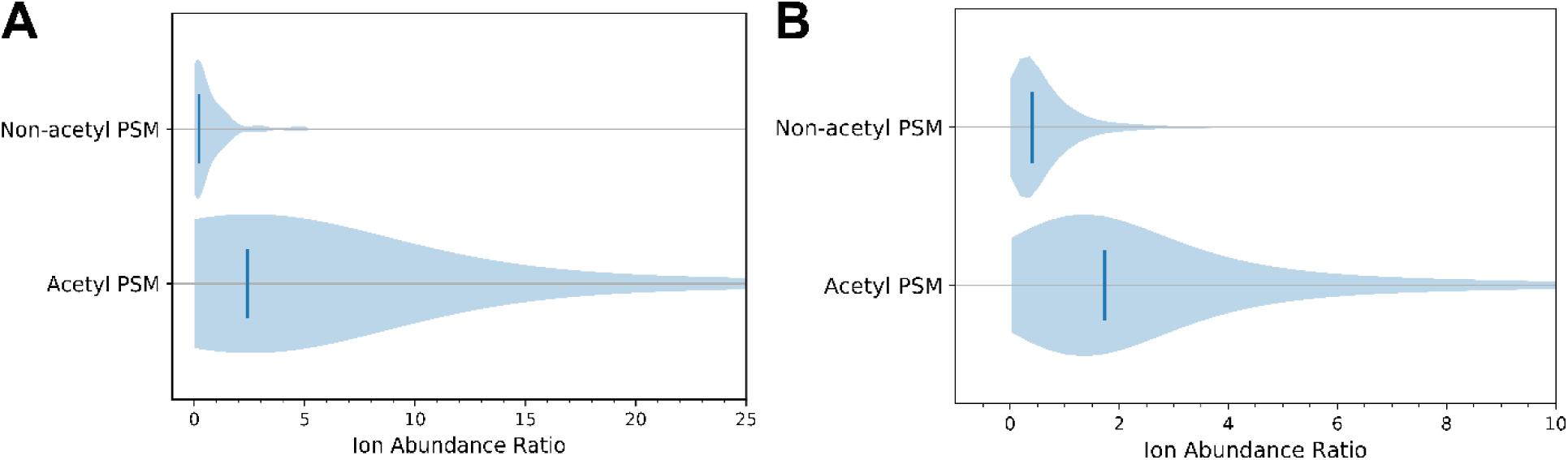
Ion abundance ratios [126]/[129] for acetylated and non-acetylated spectra. A) Violin plots of the [126]/[129] abundance ratio for all PSMs in acetylated-BSA. Vertical lines denote median ion abundance ratios 0.2 and 2.4 for non-acetyl and acetyl PSMs, respectively. B) Violin plots of the [126]/[129] ion abundance ratio for all PSMs in the Jurkat E6-1 cell dataset of Svinkina *et al*.^[33]^ Vertical lines denote the median ion abundance ratios of 0.4 and 1.7 for non-acetyl and acetyl PSMs, respectively.

Given the prevalence of the 126 and 129 ions in both acetylated and non-acetylated PSMs, we considered whether their abundance ratio could yield an improved diagnostic for acetylation. The abundance of the 129 ion was compared to that of the 126 ion. Only 13.6% of non-acetylated PSMs had a 126 ion of greater abundance than the 129 ion, while 72.8% of acetylated PSMs contained 126 ions at abundances exceeding the 129 ions. Thus, peak intensity ratios for acetylated and non-acetylated peptide spectra clearly differ, with acetyl PSMs having a [126]/[129] ratio greater than 1 (**Figure 1A**). Employing the [126]/[129] ratio to discern between non-acetylated and acetylated PSMs greatly increases the specificity as compared to the 126 ion alone. Similar trends could be ascertained from the previous report published by Svinkina, *et al*.^[33]^, with only 17.8% of non-acetylated PSMs and 71.8% of acetylated PSMs displaying the 126 ion at greater abundance than the 129 ion (**Figure 1B**). This data suggests that the trends identified here are generalizable to large datasets and can increase the reliability for assigning spectra with lysine acetylation.

For acetylated peptide PSMs, the 126 ion tended to be more abundant than other ions in the MS^2^ spectrum. The ion abundances within each spectrum containing a 126 ion were ranked; approximately 80% of acetyl PSMs had *m/z* 126 as one of the 10 most abundant ions, as compared to 16% of non-acetyl PSMs (**Figure 2A**). The trend is also true in the re-mined Jurkat dataset (**Figure 2B**). ^[33]^

**Figure 2.**
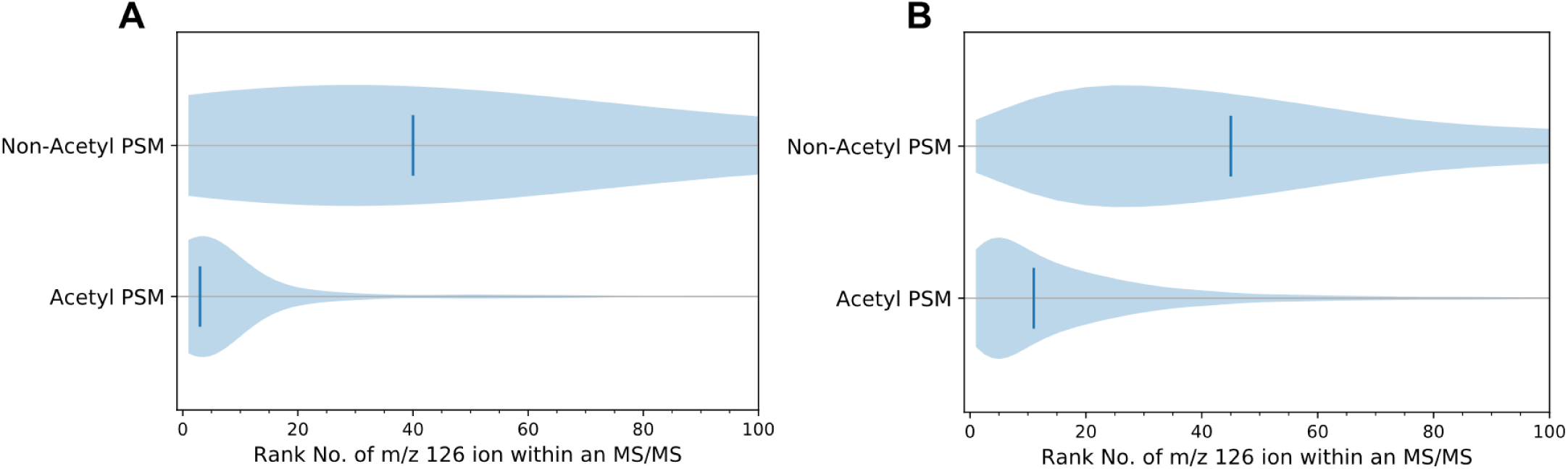
Abundance rank *m/z* 126 in MS/MS spectra of non-acetylated and acetylated peptides. A) Violin plots ranking the 126 ion abundance to all other ions in a spectrum for all acetyl-BSA PSMs. Vertical lines denote median rankings 40 and 3 for non-acetyl and acetyl peptides, respectively. B) Violin plots ranking the 126 ion abundance to all other ions present in a spectrum for all Jurkat E6-1 PSMs (dataset of Svinkina *et al*.^[33]^) Vertical lines denote median rankings 45 and 11 for non-acetylated and acetylated PSMs, respectively.

Butyrylated BSA was also analyzed in order to determine if a similar trend applied to other short-chain acylations. Data generated from butyrylated BSA had limited PSMs due to technical difficulties of the butyric anhydride used to modify the lysine side chains. Despite this, the trends were the same as acetyl BSA. The presence of the *m/*z 126 analogue for butyryl-lysine (*m/z* 154.1232) in a spectrum corresponded to a butyrylated PSM 96.4% of the time (**Table S2, Supplemental Figure S2**), and 96.2% of those spectra had *m/z* 154 as one of the 10 most abundant ions., indicating that it is both specific and sensitive.

### 3.3 Sequence Dependence of Immonium Ion Formation

Efficient formation of immonium ions can depend on a modified residue’s location in a particular peptide sequence. Sequence-specific fragmentation is a well elucidated phenomenon, as certain local residues can affect CID fragmentation; for example, when a proline or glycine residue is near.^[34–38]^ Some examples of this sequence dependence are shown in **Figure 3**. Generally an acetyl-lysine at the N-terminal position will yield the strongest signal for the 126 ion^[24]^ (**Figure 3A**). This observation can be rationalized by knowing that the first step in immonium ion formation requires an *N*-terminal amine; hence, an internal lysine would require two fragmentation events, while the *N-*terminal lysine requires only one. Sequence composition is also important. For example, two different peptides with acetylation at the K1 position yield very different relative abundances for the 126 ions (**Figure 3A**). Multiple lysine peptides that are acetylated closer to the *N-*terminus tend to yield more abundant 126 ions (**Figure 3B**). Thus, there can be a localization bias when using the cyclized immonium ion to identify and/or validate acyl modifications.

**Figure 3.**
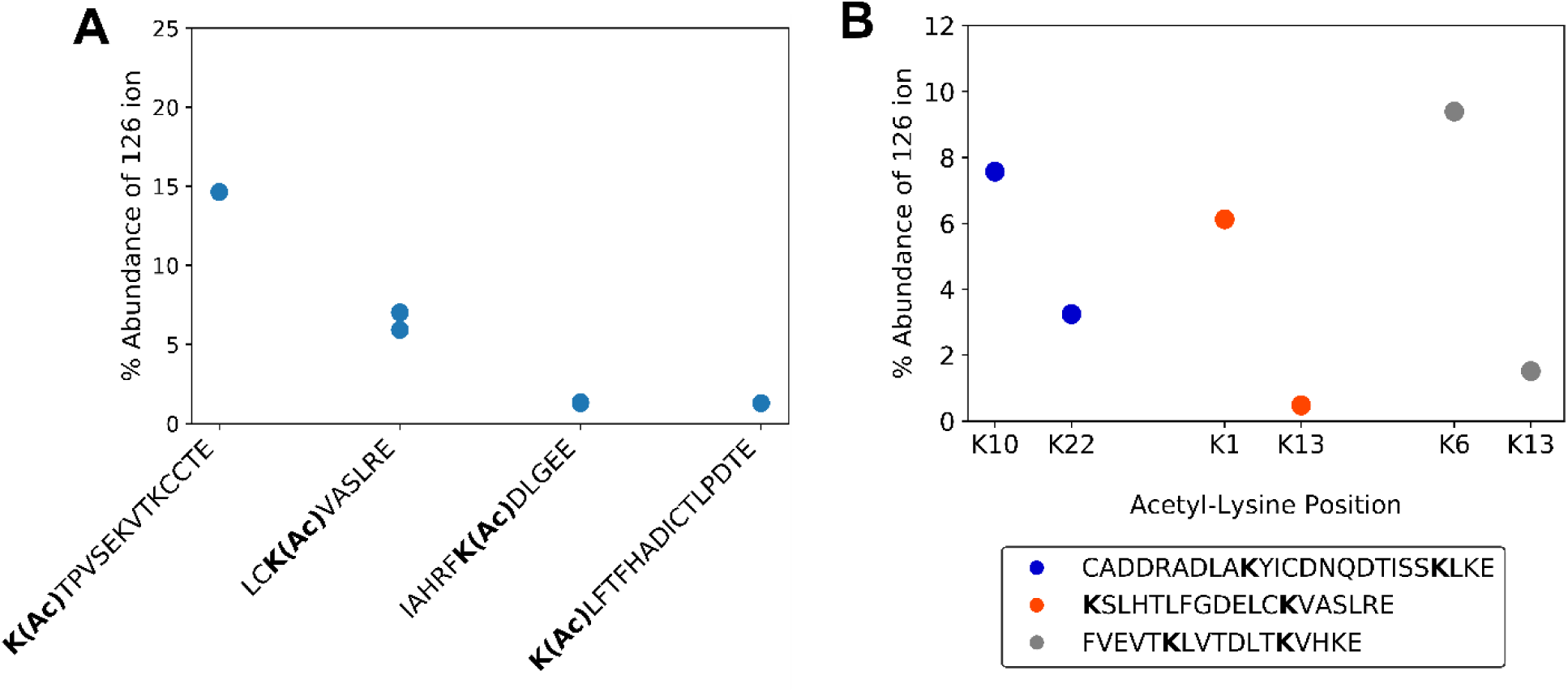
Dependence of 126 ion abundance on sequence context within fragmentation spectra. A) 126 ion abundance versus total product ion abundance for acetyl-BSA peptides. B) 126 ion abundance versus total product ion abundance for isobaric acetyl-BSA peptides. A single PSM of the same precursor charge state was selected for each acetyl-lysine position.

The normalized collisional energy (NCE) was increased in an attempt to maximize the 126 ion’s signal. Global analysis with this optimization indicates that the elevated NCE increases the relative abundance of the 126 ion from the acetylated BSA sample (**Figure 4A**). As per the violin plots, the 126 ion relative abundance distribution for PSMs increases with increasing NCE. Tandem mass spectra for individual peptides were examined to ensure that this trend applied to modifications in varied sequence locations. **Figure 4B-D** shows examples of different modified sequences. All of these peptides show that increased collisional energy increases the relative abundance of the immonium ion, congruent with the global analysis in **Figure 4A**.

**Figure 4.**
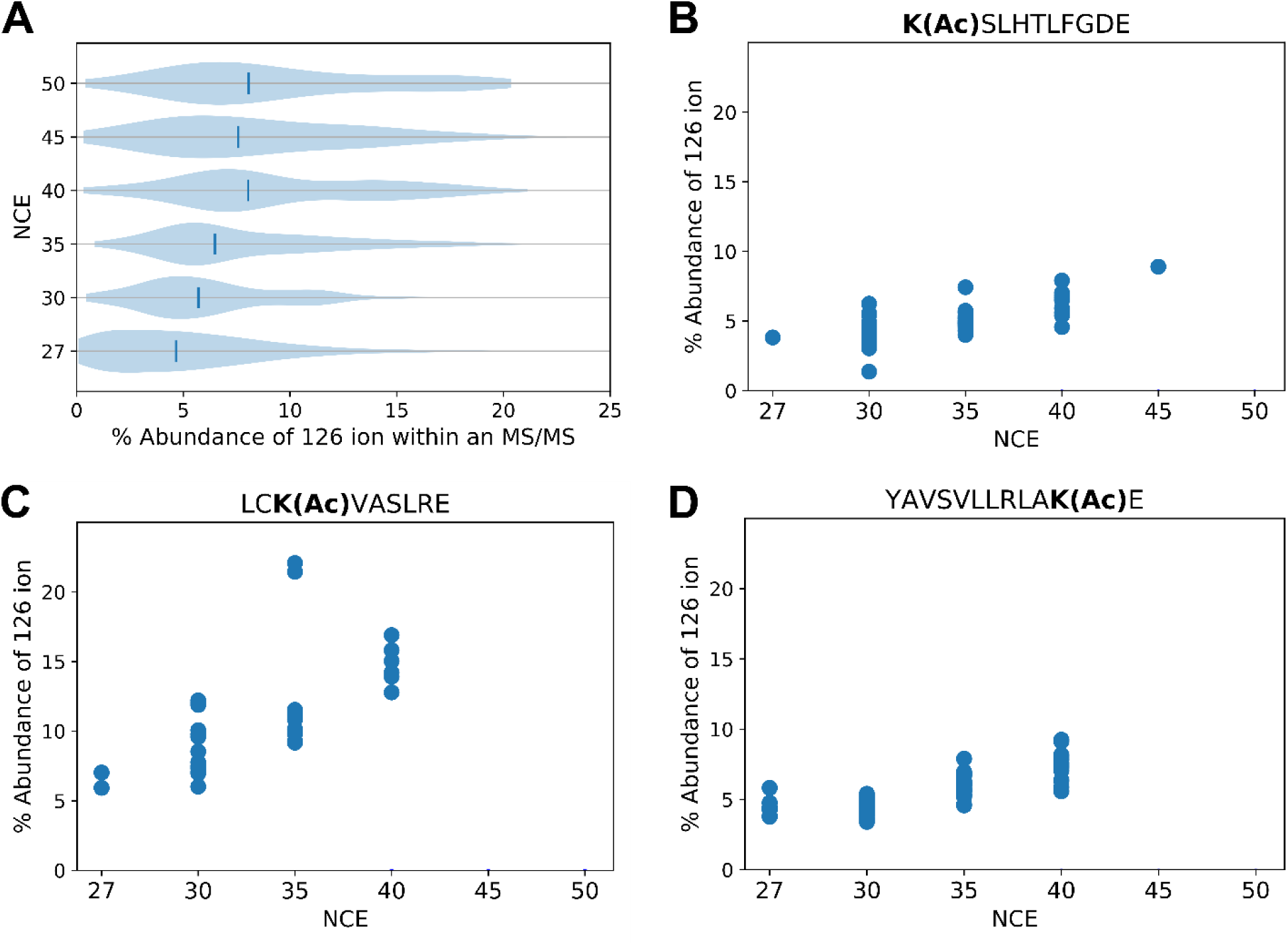
Abundance of the 126 ion versus total product ion abundance in MS^2^ spectra of acetyl PSMs. A) Each violin plot shows data obtained at a given NCE that reveals for each acetyl-PSM the percentage of total ion signal due to *m/z* 126. Vertical lines indicate the median relative abundances of the 126 ion. B-D) Percentage of total signal due to the 126 ion versus NCE for acetylated peptides selected for their varied acetyl-lysine position and sequence context. Some peptides were not identified at higher NCEs.

### 3.4 Stepped Collisional Activation Mitigates a Loss of Sequence Identifications

Increasing collision energy increases the intensities of low mass peaks, such as immonium ions, but in doing so, information critical to identifying the peptide sequence may be lost. At an NCE of 27, typical for most proteomic CID fragmentation, over 200 unique peptides could be identified from acetylated-BSA LC-MS/MS runs. At 30 NCE, the number of identified peptides dropped moderately to just below 100. However, at NCE 35 and 40, the number falls dramatically to just over 20 (**Figure 5**). While higher collision energies highlight the 126 ion in MS^2^ spectra, the benefit comes at the cost of poor sequence information, making it less likely that peptides can be matched to the correct sequence confidently.

**Figure 5.**
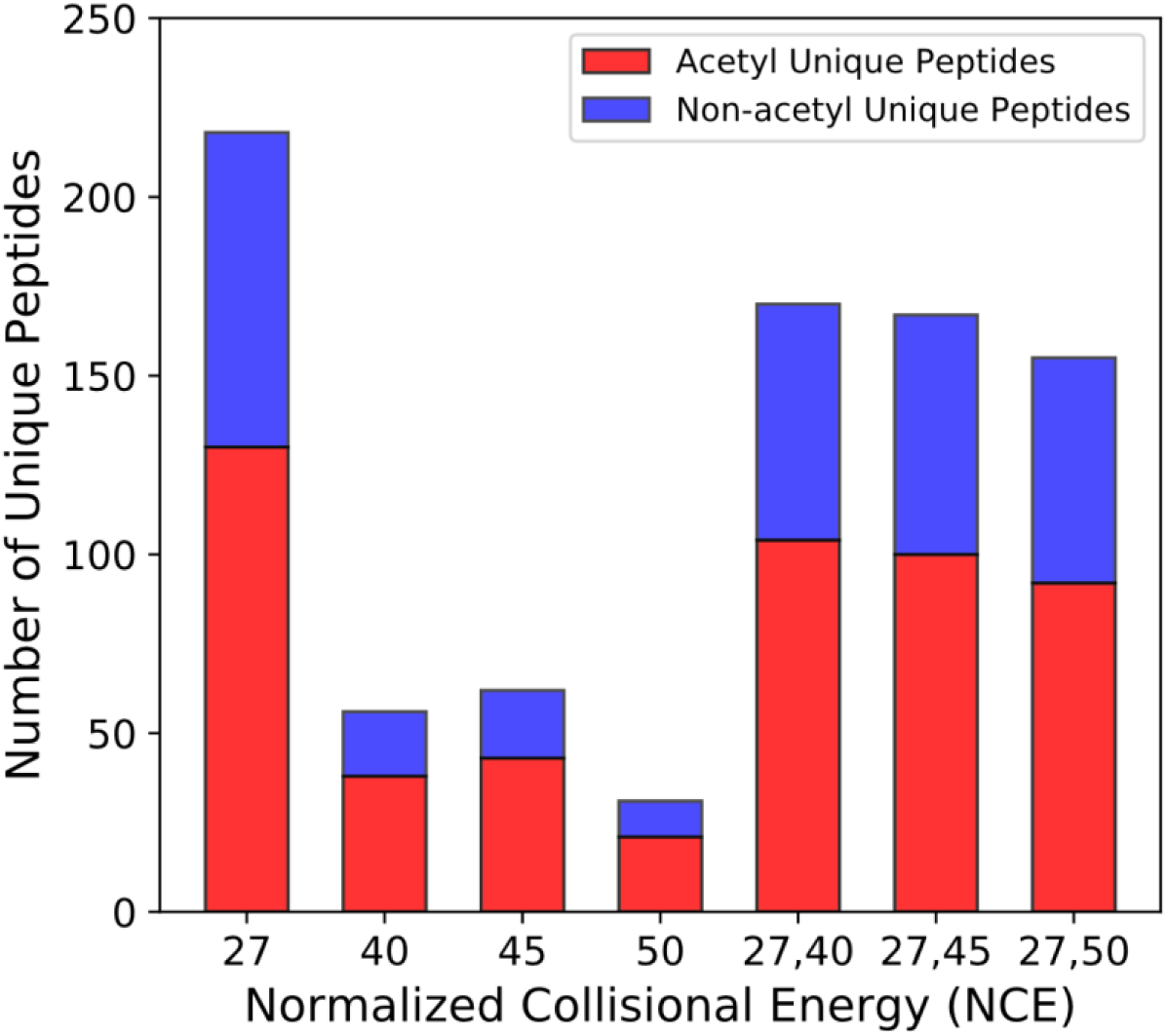
Unique peptides identified with different values of fixed and stepped NCE. Acetylated (red) and non-acetylated (blue) peptides.

To recover this information, a stepped collision energy (CE) approach was taken to enhance the intensity of the immonium ion peaks while retaining enough high mass information to identify the peptide sequence. In the stepped CE strategy, precursor ions are fragmented at low and high NCE, and the product ions from each CE are pooled for detection. **Figure 5** shows how stepped CE recovers the sequence information lost by using excessive collision energy. Several stepped collision energy combinations (NCEs of 27,40; 27,45; and 27,50) were tested to optimally balance maximal signal for the 126 ion with retention of sequence-related information. A stepped NCE method of 27,40 was found to be optimal, recovering 80% of unique peptides relative to the single energy method at 27 NCE (**Figure 5**). Applying this approach, we rescue the number of peptides identified while maintaining a strong immonium ion, such that 95.4% of acetylated PSMs present a 126 ion within the 10 most abundant ions (compared to 80% from 27 NCE). Butyrylation (*m/z* 154.1232) showed similar trends with respect to stepped NCE (**Supplementary Figure S3**).

### 3.5 Immonium Ions from Stepped Collision Energy Identify Acyl Modifications in a Complex Lysate

A question that remained with the stepped collision energy method was whether it would readily reveal immonium ions for acetyl-lysine and other acyl-lysines (*e.g.*, butyryl- and crotonyl-lysine [*m/z* 152.107]) in a complex biological sample. To investigate this question, synthetic acylated peptides were spiked into a trypsinized HeLa lysate from which the respective immonium ion: *m/z* 129 abundance ratios were monitored. The spiked peptides (**K**STPEAMAK, F**K**DEIPVVIK, STDP**K**GPSVR) differ in the type and location of acylation. In additional to acetyl-lysine, we chose butyryl- and crotonyl-lysine because they are relevant to *Syntrophus aciditrophicus*, a syntrophic bacterium for which we expect extensive protein acylation due to high cellular concentrations acyl-CoA metabolites.^[17]^ Our lab identified these sequences as highly modified in the course of *S. aciditrophicus* proteomic studies.^[39]^ Plotted are the cyclized immonium: 129 ion relative abundance ratios for each MS^2^ spectrum (**Figure 6**). Acylated peptides are clearly identified in the population of PSMs with [immonium]/[129] ratios exceeding 1. Isomers of the 126 ^[24]^ and 152 ions are also present (**Supplemental Figure S4**) and some peptides containing these sequences were identified (**Figure 6**). The quality of some MS^2^ spectra with abundance ratios exceeding 1 was insufficient to ascribe a sequence. These instances are labeled as unidentified in **Figure 6**.

**Figure 6.**
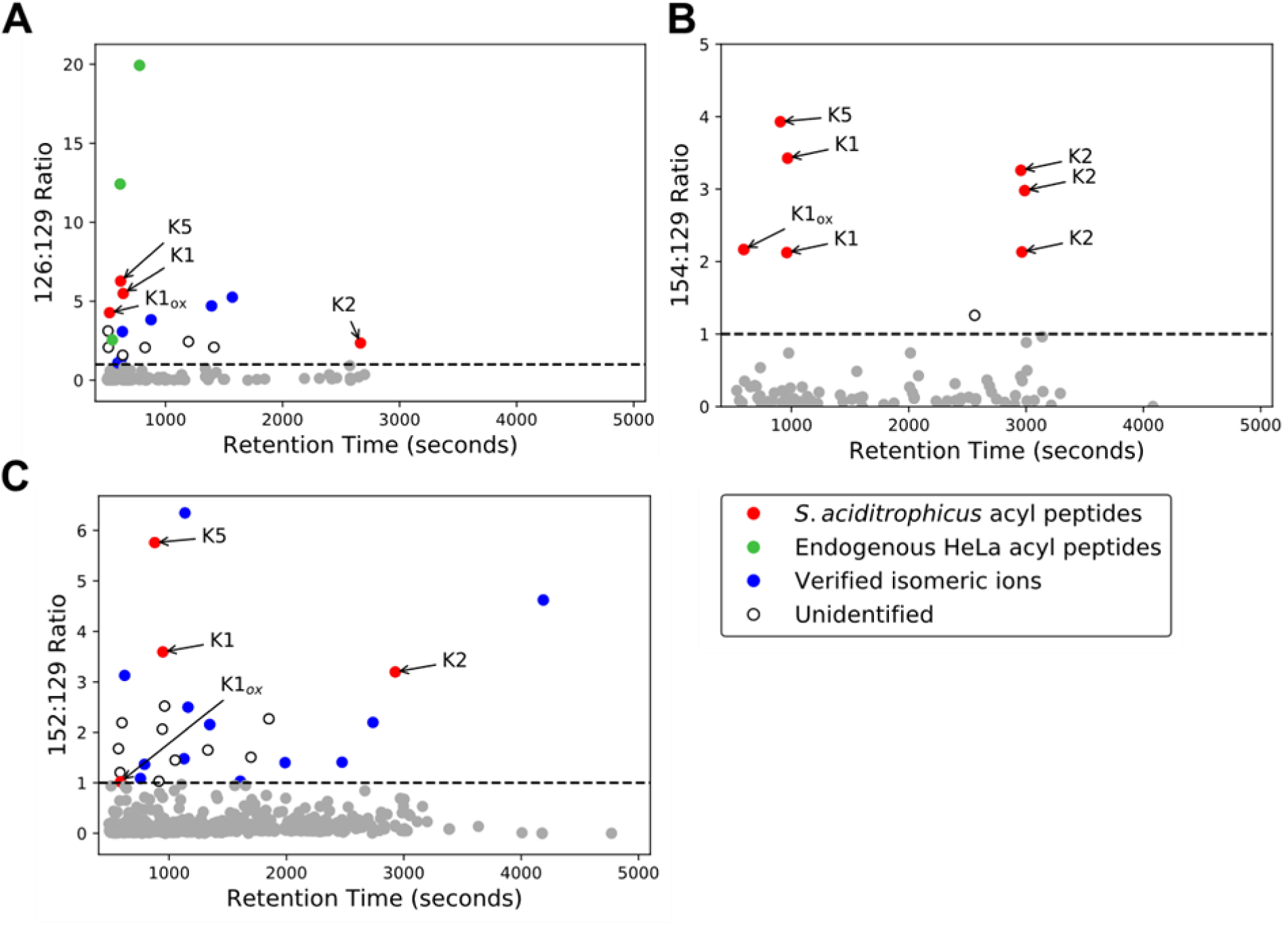
Immonium ion abundance ratio reveals low abundance synthetic acyl-peptides in a HeLa lysate background. Scatterplots of the immonium ion abundance ratio versus retention time for A) lysine acetylation ([126]/[129]), B) lysine butyrylation ([154/[129]], and C) lysine crotonylation ([152]/[129]), respectively. Grey dots represent spectra with an abundance ratio less than one. K1, K2, and K5 correspond to the synthetic *S. aciditrophicus* sequences **K**STPEAMAK, F**K**DEIPVVIK, and STDP**K**GPSVR, respectively, where the bolded residues are acylated. Verified isomeric ions and unidentified ions are described in the text.

### 3.6 Using Immonium Ions to Identify the Presence of Novel PTMs

*S. aciditrophicus* has unique metabolic pathways that allow for the formation of novel protein PTMs. Given that its short chain and aromatic fatty acid metabolism generates a variety of reactive acyl-CoA metabolites, we expect the *S. aciditrophicus* proteome to display unique acyl-lysine modifications. These acyl-CoA intermediates can spontaneously acylate lysine side chain amines under physiological conditions.^[40]^ One acyl-CoA intermediate within the benzoate degradation pathway is *3-*hydroxypimelyl-CoA.^[41]^ Interestingly, we noted that some mass spectra from the benzoate-cultivated *S. aciditrophicus* proteome presented mass shifts of K +158.0579 Da. To determine the validity of this novel modification, immonium-related ions were examined to verify whether the shift was associated with a novel lysine PTM on a benzoate-CoA ligase peptide, or if it was a misidentification.^[42]^ Based on the immonium ion structure, one expects certain facile neutral losses (**Figure 7**) to be present, an NH_3_ neutral loss (similar to that of the acetyl-lysine 126 ion), an H_2_O loss, and the loss of both NH_3_ and H_2_O. Tandem mass spectra from two peptides containing the putative modification, benzoate-CoA ligase (RS03815/RS03820) and acetyl-CoA transferase (RS12490), presented ions corresponding to neutral loss of NH_3_, H_2_O, and NH_3_+H_2_O from the predicted immonium ion (**Figure 7, Supplemental Figure S5**); *i.e.*, 242.1392, 241.1552, and 224.1278 Da, respectively. These immonium-related ions validated the presence of a hydroxypimelyl-lysine, the first observation of this PTM.

**Figure 7.**
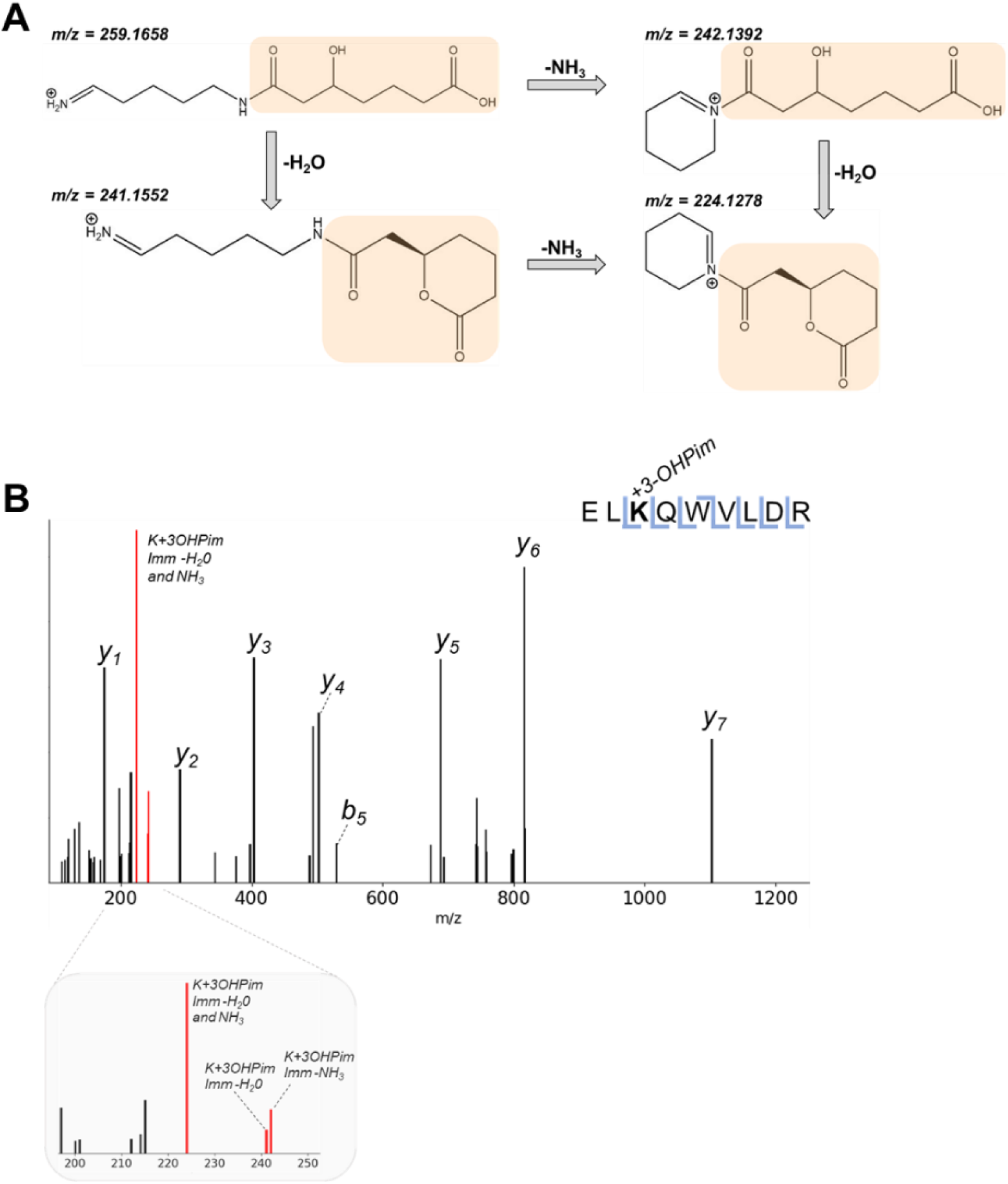
Novel modification (3-hydroxypimelylation) on benzoate-CoA ligase in *S. aciditrophicus* with its diagnostic ion. A) Immonium ion for for 3-hydroxypimelylation (K+158.0579 Da) (highlighted in orange) undergoes facile neutral losses. B) MS/MS spectrum showing mass shift corresponding to 3-hydroxypimelylation and its low mass region enlarged to highlight immonium-related ions (colored in red).

## 4. Discussion

Immonium and immonium-related ions are excellent proxies for the presence of acetylation.^[24,43]^ Despite their long use in global analyses, current instrumentation and methods have not been optimized for their use. While previous reports found that the specificity for acetylation of the *m/z* 126 ion was superior to that of m/z 143,^[24]^ the former is often observed in mass spectra lacking any acetylation, as is confirmed in our analysis. Hence, the diagnostic ion’s presence alone is not sufficient to claim modification with high confidence, or to restrict data acquisition or analysis to only acetylated peptides. Incorporating abundance ratios provides a nuanced way to utilize this marker. Likewise, incorporating a stepped collisional energy CID experiment may increase confidence in acyl-lysine assignments. Akin to oxonium ion analysis, using a stepped NCE strategy highlights the diagnostic ions of acyl-lysine while also optimizing sequence-related information in a data dependent acquisition (DDA) method. Stepped NCE is particularly important for modifications present in the middle of a peptide, as generating these immonium ions require more collision energy.

Future applications of stepped NCE with immonium ion abundance analyses can take two possible routes. One is to expose and verify novel PTMs. As the list of putative acyl-modifications grows,^[4,44,45]^ it becomes critical to obtain additional constraints (metrics) to validate the presence of increasingly complex modifications. Many acyl-modified peptides can be isobaric with di- or tri-peptide related ions or be indistinguishable from non-acylated ions in low resolution instruments.^[42]^ For example, acetyl and propionyl modifications differ by a single methyl group, as do propionyl and butyryl modifications. The presence of a methylated residue or an amino acid that differs from another by a methyl group (Ser/Thr, Asn/Gln, Ala/Val, etc) may make it impossible to distinguish the correct acyl modification.^[46]^ Similarly, carbamylation has a 43.00582 Da mass shift, whereas acetylation combined with deamidation produce a 42.99458 Da shift. Depending on the mass of the ion and the resolving power of the instrument, these PTMs could be indistinguishable in the event of poor fragmentation and unincorporated diagnostic ions.

Stepped NCE can also apply in conjunction with data independent acquisition (DIA) approaches. Currently, DIA is greatly limited by the complexity of spectral deconvolution, particularly by fragment ion interference when concurrently eluting peptides have very similar masses.^[47]^ Diagnostic ions, however, can increase the confidence that an acyl-lysine precursor is present in the convoluted spectra; thereby providing more information for elegant algorithms to consider when identifying and assigning PTMs. Furthermore, experiments targeting acyl modifications can exploit immonium and immonium-related ions in parallel reaction monitoring (PRM) or product ion scanning.

Increased understanding of acyl modifications has suggested some biological functions for these modifications.^[48]^ Given the reactive nature of acyl-CoA species^[17]^ and the new depths proteomics can reach, it should be expected that more acyl modifications will be found and may show biological significance in metabolic pathways across organisms. As datasets grow in size and complexity, confident assignments are essential. Properly incorporating immonium ions into the assignment of acyl-modifications, will increase confidence in established PTM identifications and support/validate assignment of those yet to be discovered.

## Acknowledgements

Funding from the Department of Energy Office of Science (BER) contract DE-FC-02-02ER63421 (to J.A.L.; UCLA/DOE Institute for Genomics and Proteomics), NIH Ruth L. Kirschstein National Research Service Award (to J.Y.F.; GM007185), and NSF Graduate Research Fellowship (to J.Y.F.; DGE-1650604) is acknowledged. J.M.M. was supported by a UCLA Molecular Biology Institute Whitcome Fellowship. We would also like to thank Drs. Michael J. McInerny and Robert P. Gunsalus for the *S. aciditrophicus* samples used in the study.

## Supplemental Tables and Figures

**Table S1.** Distribution of PSMs containing the *m/z* 101 immonium ion from acetylated BSA (27 NCE) and in Jurkat E6-1 cells (25 NCE) (Svinkina *et al.*) ^[33]^

|  |  | PSMs with<br>101 ion |  | PSMs without<br>101 ion |  |
| --- | --- | --- | --- | --- | --- |
| Acetyl-BSA | Non-acetyl | 59 | (25%) | 177 | (75%) |
|  | Acetyl | 58 | (17.3%) | 277 | (82.7%) |
| Svinkina et al. | Non-acetyl | 201 | (3.3%) | 5834 | (96.7%) |
|  | Acetyl | 151 | (3.3%) | 4449 | (96.7%) |

**Figure S1.**
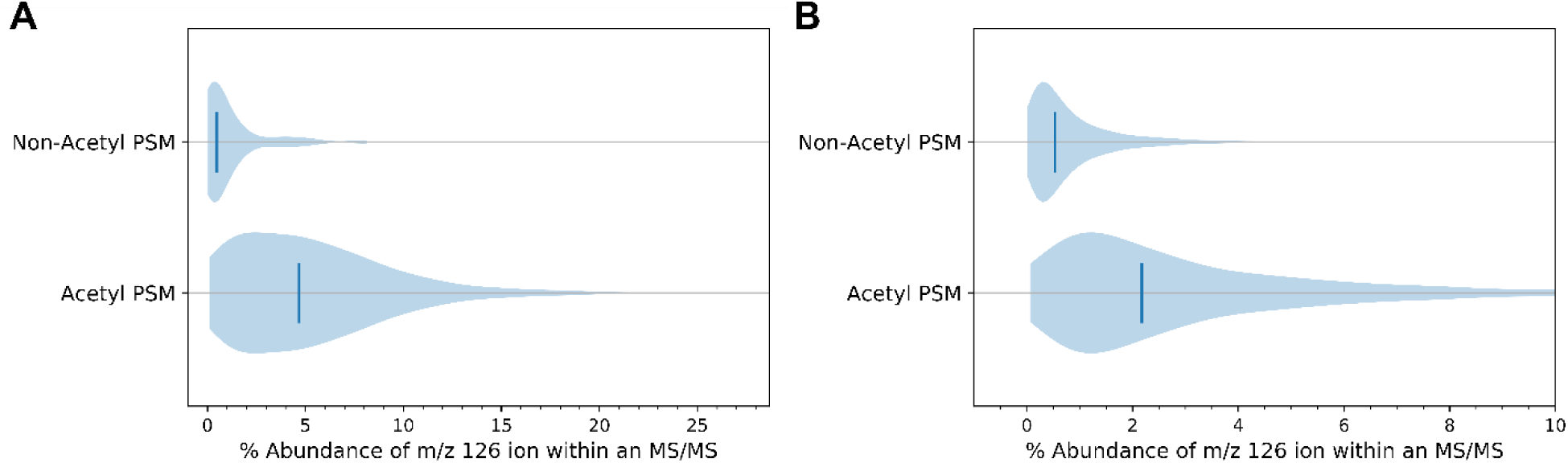
Percentage of total MS^2^ ion signal for *m/z* 126. A) Violin plots of the relative signal from *m/z* 126 for acetylated and non-acetylated PSMs of acetyl-BSA. Vertical lines denote the 0.5% and 4.7% median percent intensity from non-acetyl and acetyl PSMs, respectively. B) Violin plots of the relative signal from *m/z* 126 for all PSMs in the Jurkat E6-1 cell dataset of Svinkina *et al.*^[33]^ Vertical lines denote the 0.5% and 2.2% medians for non-acetyl and acetyl PSMs, respectively.

**Table S2.** Distribution of PSMs containing 154 and 129 immonium ions in peptides from butyrylated BSA (27 NCE).

|  |  | **PSMs with**<br>**154 ion** |  | **PSMs without**<br>**154 ion** |  | **PSMs with**<br>**129 ion** |  | **PSMs without**<br>**129 ion** |  |
| --- | --- | --- | --- | --- | --- | --- | --- | --- | --- |
| Butyryl-BSA | Non-butyryl | 4 | (18.2%) | 18 | (81.8%) | 15 | (68.2%) | 7 | (31.8%) |
|  | Butyryl | 27 | (96.4%) | 1 | (3.7%) | 16 | (57.1%) | 12 | (42.9%) |

**Figure S2.**
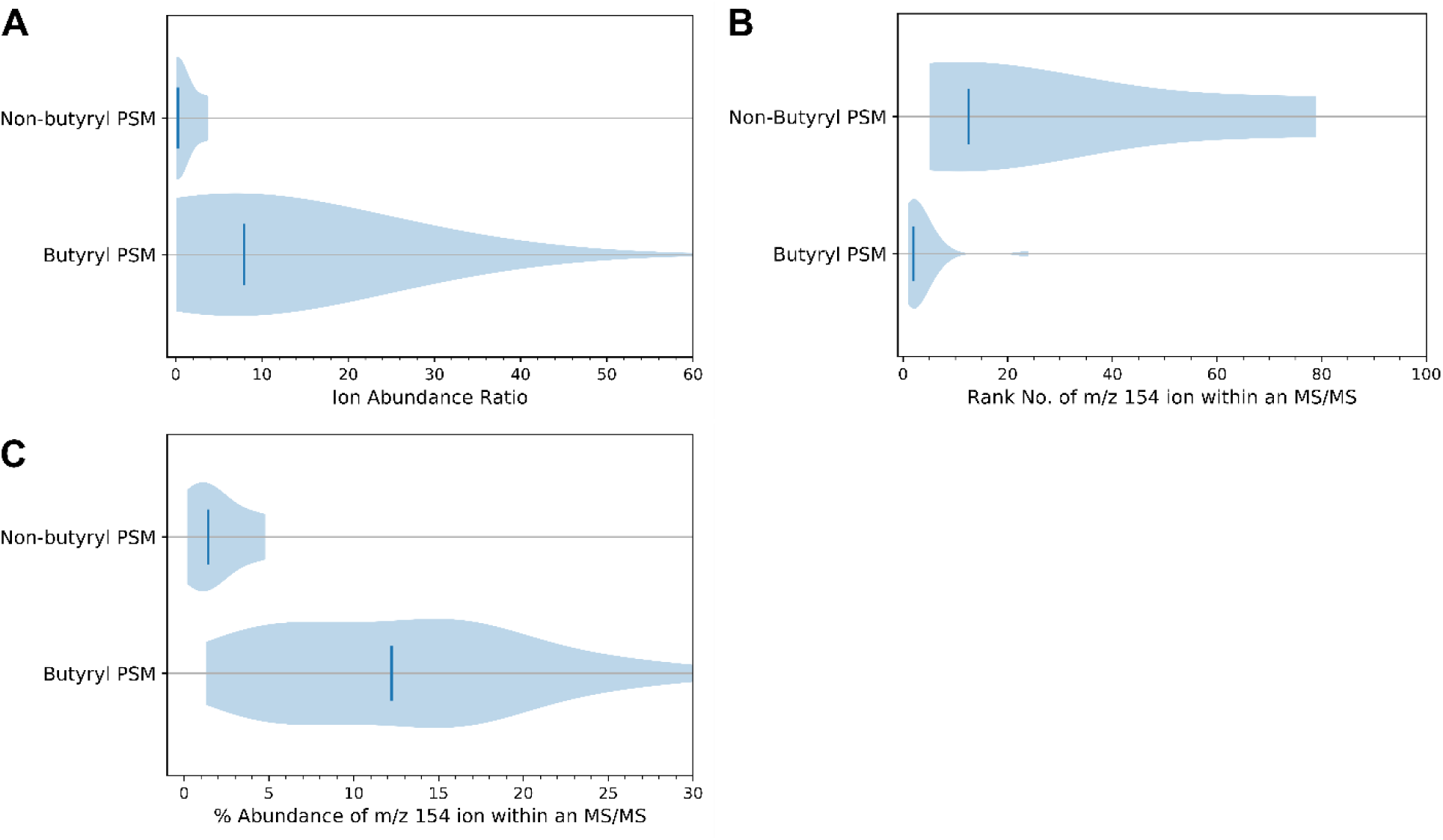
The 154 ion and [154]/[129] abundance ratios as markers for butyryl-lysine at 27 NCE. A) Violin plots of the [154]/[129] ion abundance ratio for all butyryl-BSA PSMs. Vertical lines denote the, 0.3 and and 8.0 median ion abundance ratios for non-butyryl and butyryl PSMs, respectively. B) The abundance rank of *m/z* 154 relative to other product ions in each spectrum. Vertical lines denote the median ranks, 13 and 2, for non-butyryl and butyryl PSMs, respectively. C) The percentage of total product ion signal from *m/z* 154. The vertical lines denote the median percent abundance, 1.4% and 12.2%, for non-butyryl and butyryl PSMs, respectively.

**Figure S3.**
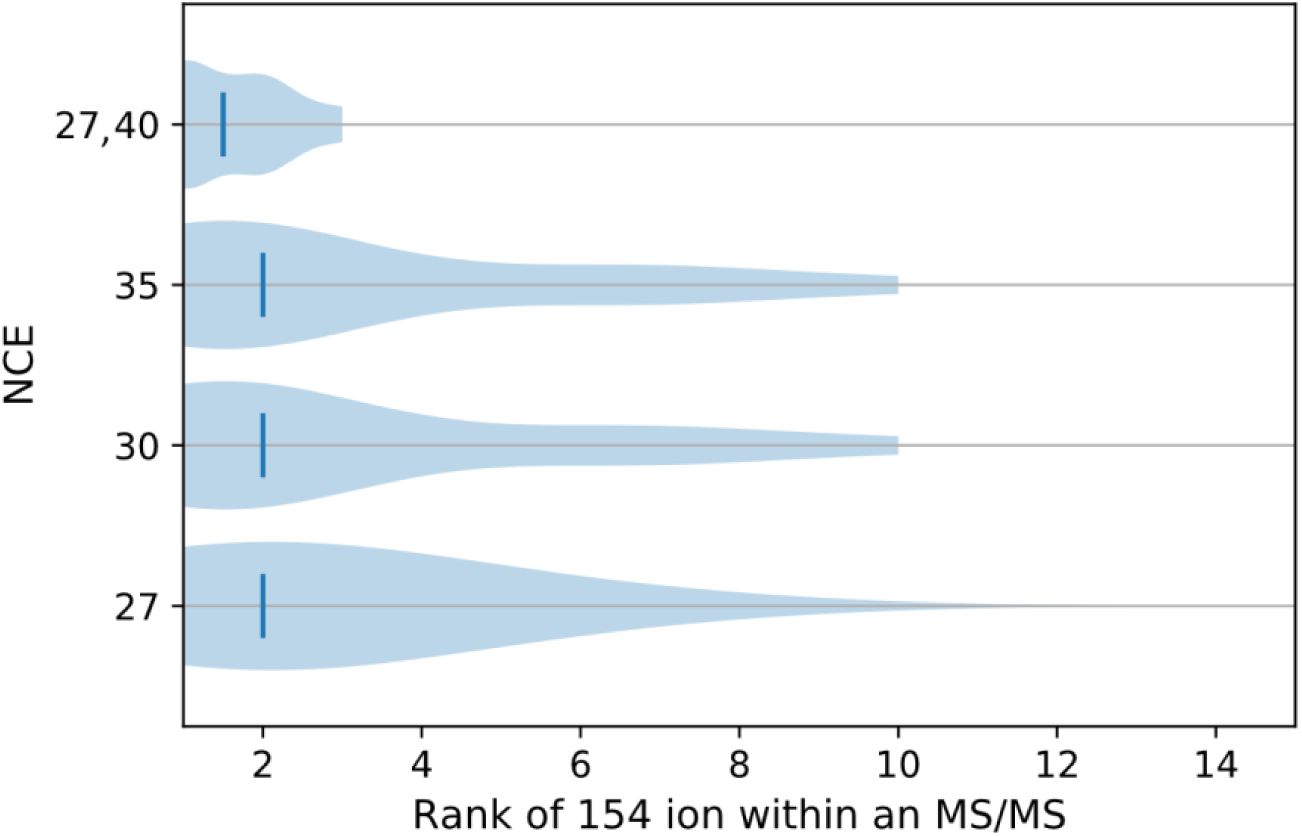
Abundance rank of the 154 ion depends on NCE setting. Violin plots of the 154 ion’s abundance rank in a spectrum for butyryl PSMs. Vertical lines denote the median rankings: 2 for fixed NCE= 27, 30, and 35 and 1 for stepped NCE= 27,40. No butyryl PSMs were identified at 40, 45, and 50 NCE.

**Figure S4.**
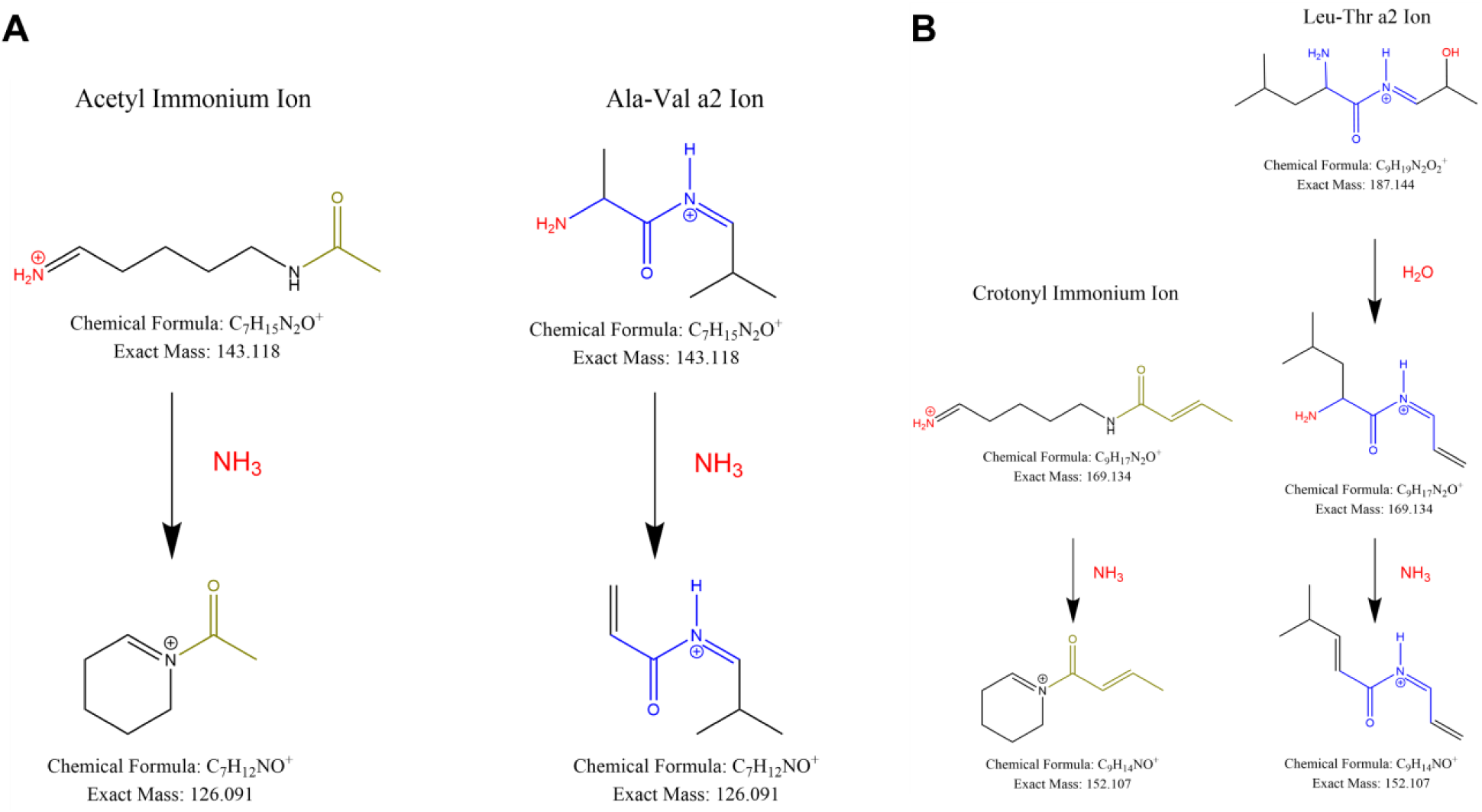
Verified isomers of acyl-lysine marker ions. Proposed ions that are isobaric to the acyl-lysine immonium ions seen in abundance in Figure 6. A) Pathway producing Ala-Val products isobaric with the acetyl-lysine immonium ion. Not shown are Xle-Gly products that are also isobaric. B) Pathway producing Xle-Thr products isobaric with the crotonyl-lysine immonium ion.

**Figure S5.**
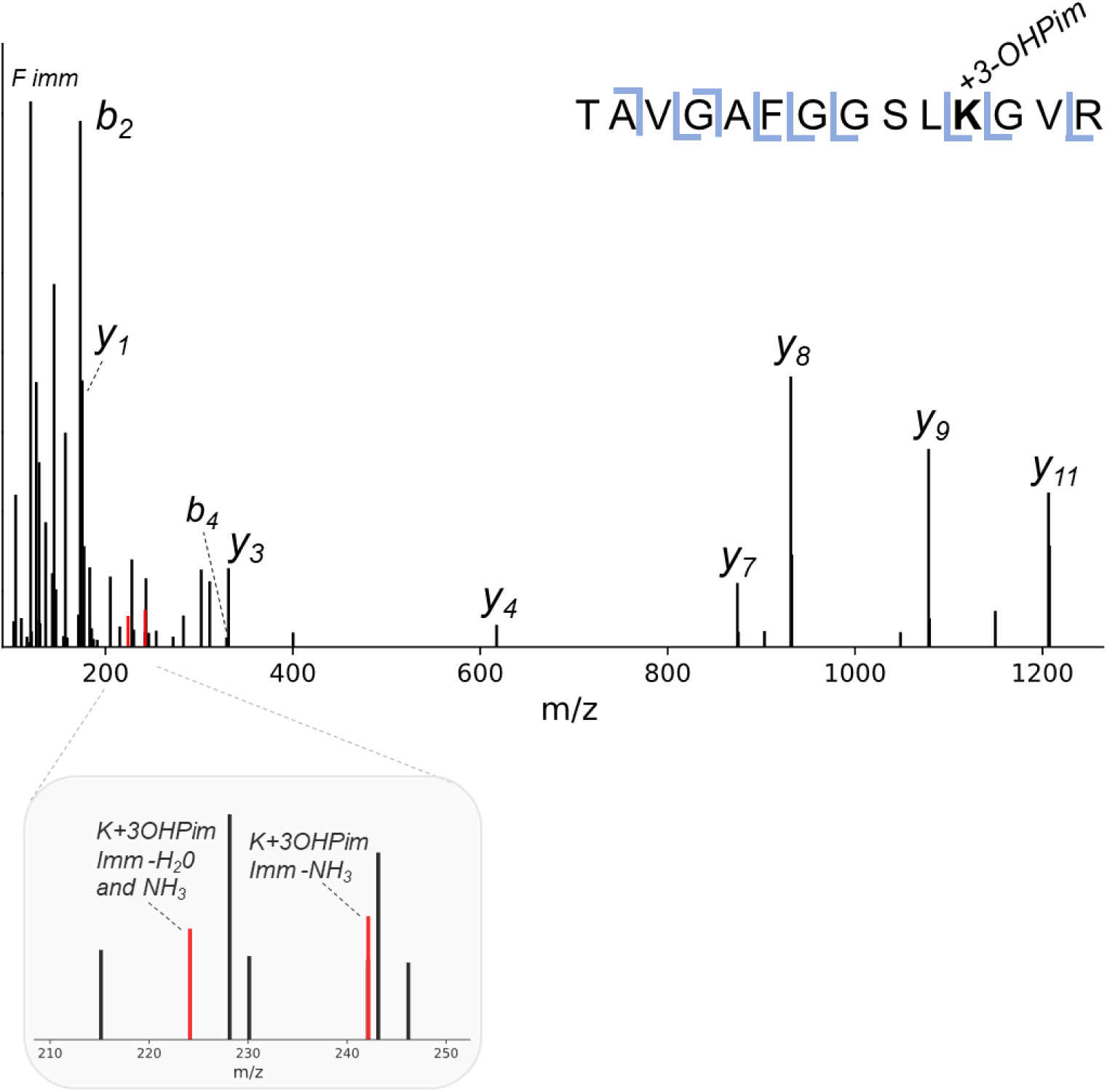
Novel modification (3-hydroxypimelylation) on acetyl-CoA acetyltransferase in *S. aciditrophicus* with its diagnostic ion. MS/MS spectrum showing mass shift corresponding to 3-hydroxypimelylation and its low mass region enlarged to highlight immonium-related ions (colored in red).

